# Serum-mediated cleavage of *Bacillus anthracis* Protective Antigen is a two-step process that involves a serum carboxypeptidase

**DOI:** 10.1101/267369

**Authors:** David L. Goldman, Edward Nieves, Antonio Nakouzi, Johanna Rivera, Ei Ei Phyu, Than Htut Win, Jacqueline Achkar, Arturo Casadevall

## Abstract

Much our understanding of the activity of anthrax toxin is based on *in-vitro* systems, which delineate the interaction between *B. anthracis* toxins and the cell surface. These systems however, fail to account for the intimate association of *B. anthracis* with the circulatory system, including the contribution of serum proteins to the host response and processing of anthrax toxins. Using variety immunologic techniques to inhibit serum processing of *B. anthracis* Protective Antigen (PA) along with mass spectrometry analysis, we demonstrate that serum digests PA via 2 distinct reactions. In the first reaction, serum cleaves PA_83_ into 2 fragments to produce PA_63_ and PA_20_ fragments, similar to that observed following furin digestion. This is followed by carboxypeptidase-mediated removal of the carboxy-terminal arginine and lysine residues from PA_20_.

## Importance

Our findings identify a serum-mediated modification of PA_20_ that has not been previously described. These observations further imply that the processing of PA is more complex than currently thought. Additional study is needed to define the contribution of serum processing of PA to the host response and individual susceptibility to anthrax.

## INTRODUCTION

*Bacillus anthracis* is the causative agent of anthrax and is widely recognized for its potential use as an agent of bioterrorism. *B. anthracis* secretes 2 bipartite toxins, lethal and edema toxins that are essential for virulence. Both toxins require the protective antigen (PA) component to mediate cell entry. PA is, therefore, essential to the damaging effects of anthrax toxins and PA-deficient mutants have significantly reduced virulence (1).

The current paradigm of toxin pathogenesis posits that *B. anthracis* secretes the pro-form of PA (PA_83_), which binds to cell surface receptors (tumor endothelium marker-8 or capillary morphogenesis protein -2 where it undergoes cleavage by cell-associated furin into 2 fragments, PA_20_ and PA_63_. PA_63_ subsequently undergoes heptamerization to form a pre-pore structure that binds edema factor (EF) or lethal factor (LF) and is internalized. Understanding the mechanism by which anthrax toxin is processed is important because interference with the processing steps is the basis for the development of therapeutics including furin inhibitors (2). In addition, antibodies reactive to PA are protective in animal models of anthrax and one monoclonal antibody, Raxibacumab, has been licensed for clinical use (3–5).

Much of our understanding about toxin processing in anthrax pathogenesis is based on experiments using in-vitro systems (Reviewed in (6)). These systems generally do not take into account the role of host serum proteins as part of the host response to anthrax. During the course of anthrax, *B. anthracis* encounters serum proteins at multiple stages including invasion into the lymphatic system and high-level bacteremia, which occurs in the context of sepsis. In late stages of experimental anthrax in macaques, for example, lethal toxin concentrations on the order of 10 μg/ ml have been reported (7). The intimate association between *B. anthracis* and serum is further highlighted by the presence of pathogen-associated proteins that directly act on elements within circulation. This includes enzymes that digest host hemoglobin and circulating lethal toxin, which interferes with neutrophil function.(8, 9).

Several lines of evidence suggest more complexity to PA processing that is apparent from the current model. Anthrax toxin is released from *B. anthracis* in vesicles that contain all toxin components (10). Although these vesicles may be rapidly disrupted by serum albumin releasing toxin components (11), they are also released intracellularly. In addition, PA **circulating in the serum** is found in animal models as a complex of PA_63_ bound to LF or EF, not as intact PA_83_ (12). In fact, serum from humans and other species has proteolytic activity that digests PA in a manner similar to furin (13–15). Our previous studies suggest a correlation between, serum-mediated digestion of PA and protection from the killing effects of Lethal Toxin *in vitro* (15). In the current work, we find that serum-mediated processing of PA is a 2-step reaction that involves carboxypeptidase-mediated truncation of the PA_20_ fragment.

## RESULTS

### Serum-mediated digestion of rPA

Serum treatment of rPA_83_ produced 2 protein fragments, PA63 and a band that is slightly lower in molecular mass than PA_20_ (Figure 1; lane 6). The larger protein is similar in size to the PA63 produced by furin digestion of rPA_83_. However the smaller protein is smaller than the PA_20_ produced by furin digestion of rPA_83_ and is referred to as truncated PA_20_. Furthermore, serum treatment of rPA_83_ before or after furin digestion still produced this truncated fragment (Figure 1, lanes 2 and 4). Heat inactivation of serum prevented this truncation (Figure 1; lanes 3 and 5), consistent with the idea that the enzyme responsible for truncation is heat labile.

**Figure 1.**
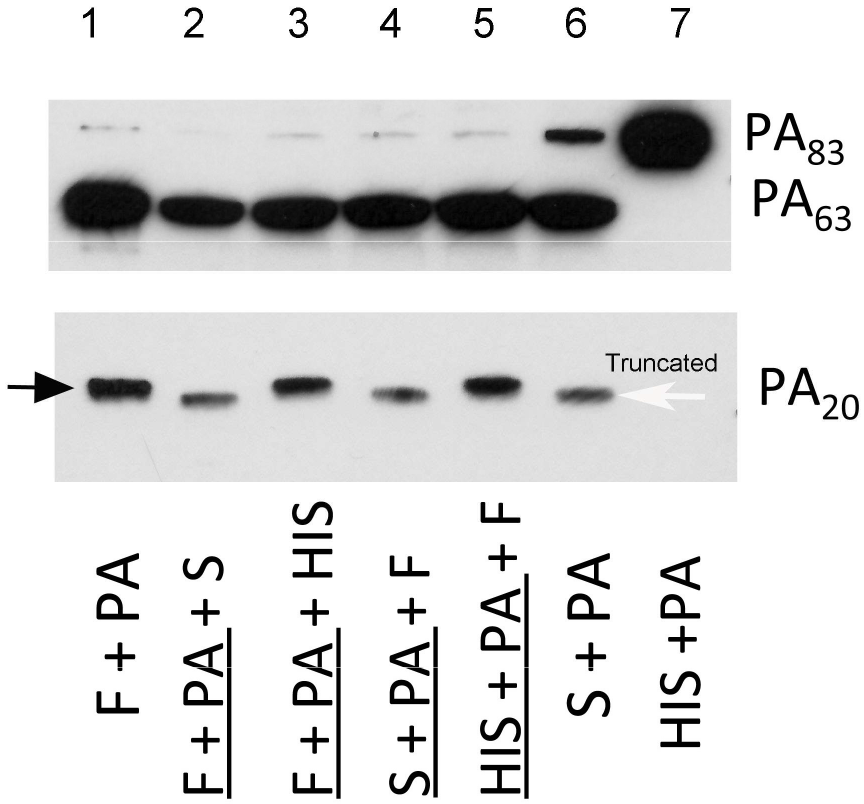
Serum mediated digestion of rPA_83_ produces a truncated PA_20_ fragment when compared with furin-mediated digestion. Shown are the digestion fragments of PA_83_ when incubated with either furin (F, lane 1), serum (S, lane 6) or heat-inactivated serum (HIS, lane 7). Treatment of PA_83_with serum either after or prior to furin digestion (lanes 2 and 4 respectively) produced a truncated PA_20_ fragment indicating that serum digestion of PA_20_ occurs with furin-digested PA. In contrast, incubation of furin-treated PA_83_ with heat-inactivated serum (lanes 3 and 5) did not produce a truncated PA_20_ fragment. For the purpose of this assay, mAb 10F4 (which recognizes domain 2-4) was used to detect the PA_63_ fragment, while mAb 19D9 (which recognizes domain 1) was used to detect both the normal and truncated PA_20_ fragments. Black arrow points to the normal PA_20_ fragment while the white arrow points to the truncated PA_20_.

### Inhibition of serum-mediated digestion of rPA

To determine the precise site at which serum cleaves rPA, we attempted to inhibit serum -mediated cleavage using a library of overlapping peptides, which represent the PA sequence and antibodies that recognize various PA sites. Pre-incubation of rPA with the mAb 19D2, which recognizes an epitope immediately C-terminal of the furin site (16), prevented rPA digestion by serum and furin. This inhibition of digestion was not seen with other PA-specific antibodies, including 7.5 G, which recognizes domain 1 of PA_83_. Serum-mediated PA cleavage was also prevented by coincubation of serum with 3 overlapping peptides (D5-D7), which contain the furin digestion site, but not with other peptides (including, D12, E1, E2, which represent PA sequences approximately 30 AA residues C terminal to the furin site) (not shown).

Using chemical inhibitors while measuring PA_63_ formation, we found that the serine/cysteine protease antipain partially inhibited the formation of PA_63_. In contrast, none of the other tested protease inhibitors, including bestatin, chymostatin, E-64, leupeptin, pepstatin, phosphoramindon, pefabloc SC and aprotinin prevented PA_63_ formation. As in previous studies, we found that EDTA was a potent inhibitor of serum mediated digestion of PA_83_. By contrast, both competitive inhibitors of furin (I and II) prevented serum mediated digestion of PA. For the furin inhibitor I, concentrations as low as 0.001 mg/ml resulted in complete inhibition of serum digestion, whereas for the furin inhibitor II concentrations as low as 0.010 mg/ml produced complete inhibition of digestion (Figure 2).

**Figure 2.**
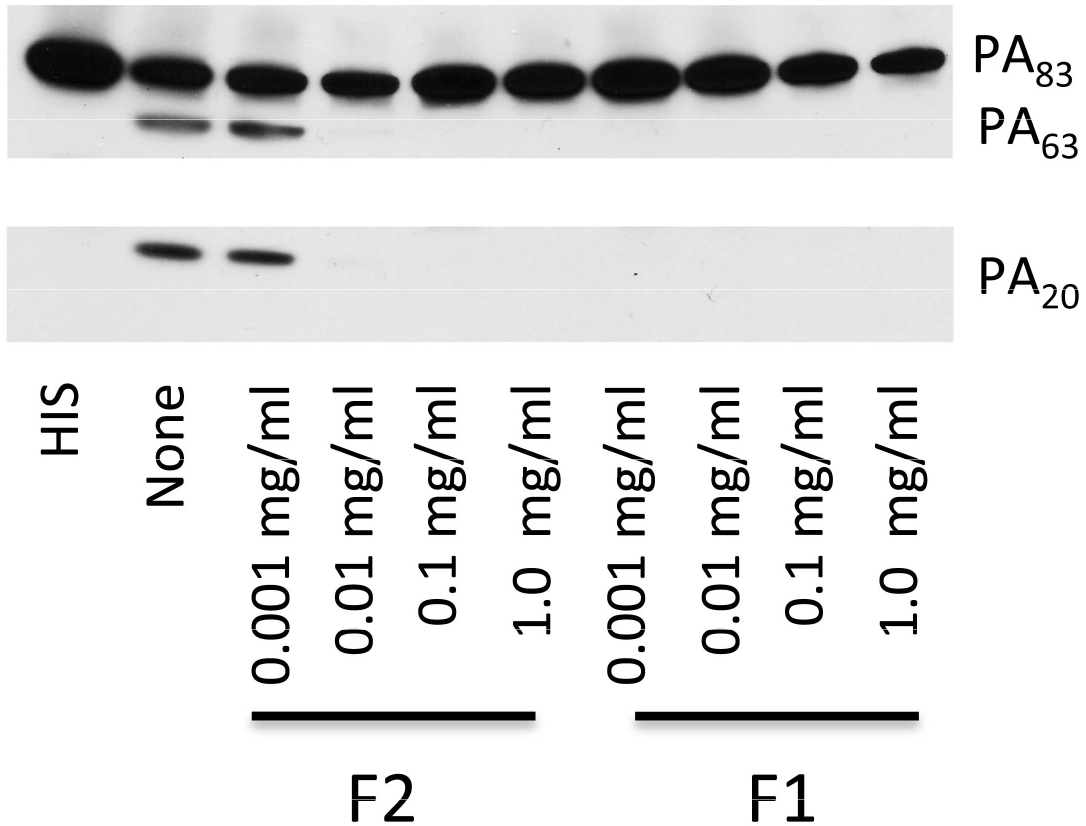
Furin inhibitors I and II prevent serum-mediated digestion of rPA_83_. Heat-inactivated serum(HIS) had no effect onrPA. In the absence of inhibitors (none) PA_83_, PA_33_ and PA_20_ -like fragment are present. Both Furin Inhibitors I and II (F_1_ and F_2_) prevented serum digestion of PA_83_. PA_83_ was incubated for serum for 30 minutes

### Truncated PA_20_ fragment

To better identify the precise site of serum-mediated digestion of rPA, the truncated PA_20_ fragment produced by serum digestion was examined by mass spectrometry. First the intact protein mass of this fragment was measured and the experimental mass determined by LC-ESI MS to be 23,600 Da (Figure 1). Furin cleaves at RXK/RR, which would correspond to a predicted molecular mass of 25,157 Da for rPA (Figure 3; n-terminus to RKKR) a difference of 1,554 Da; way beyond the error of measurement. To determine the sequence of the truncated PA_20_ fragment in-gel trypsin digestion was performed. The LC-MS/MS data identified the underlined tryptic peptides shown in Figure 3 (identified tryptic peptides of the are listed in supplemental table 1). The peptide sequence, LLNES….GFIK, is too large for fragmentation on the LTQ mass spectrometer and was not detected by MS/MS but the +4, +5, +6, +7 and +8 charge states were detected (Supplemental Figure 2). The predicted protein mass from the n-terminus to the last tryptic peptide identified is 23,213 Da and if the next 4 amino acids are included (SSNS) the predicted protein mass will increase to 23,588 Da, a difference of 12 Da or 0.05% when compared with the experimental intact protein mass (23,600 Da). These findings are consistent with serum-mediated cleavage of the basic, C-terminal arginine and lysine residues from the PA_20_ fragment produced by furin digestion followed by possibly carboxypeptidase.

**Figure 3.**
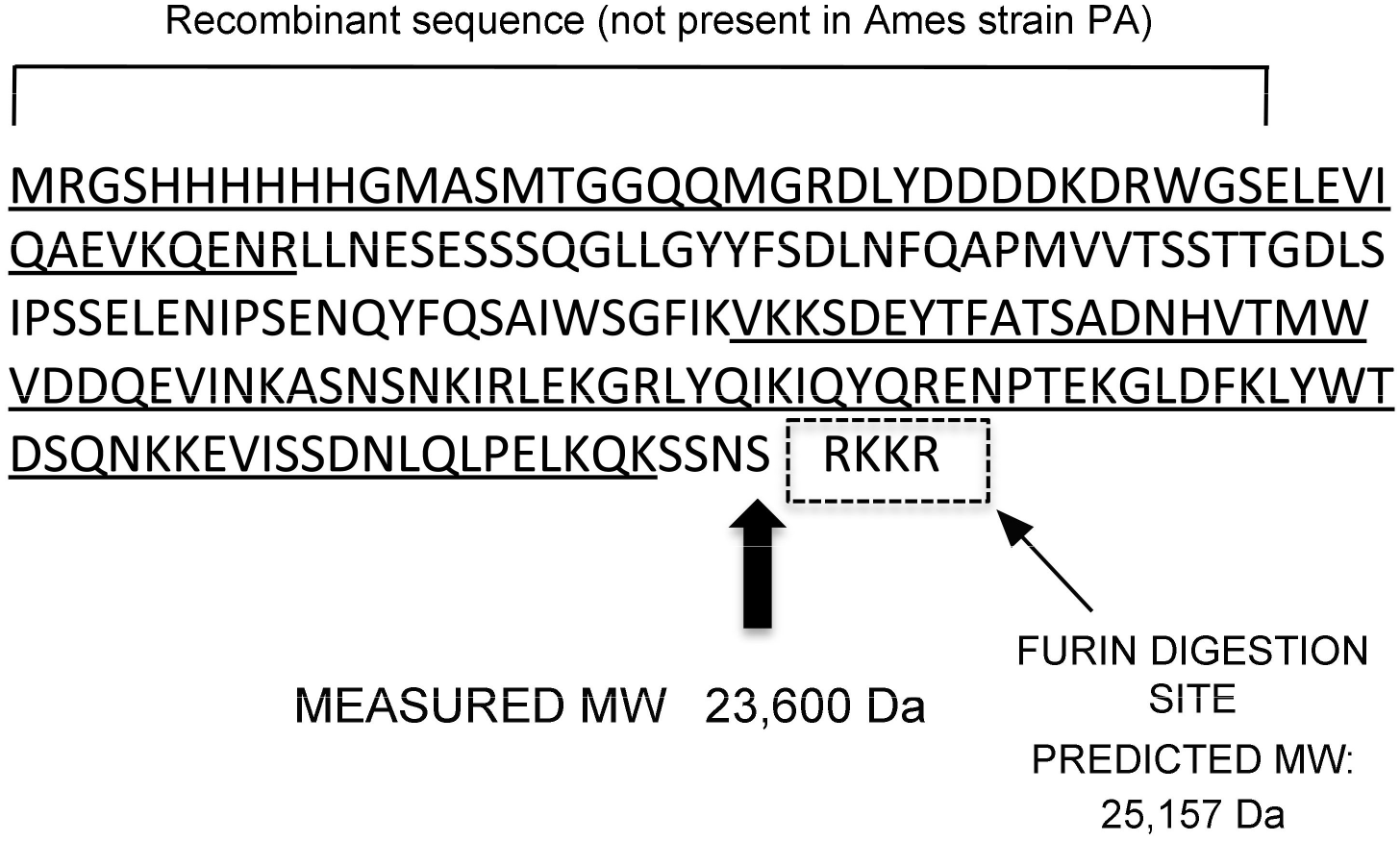
Mass spectrometry of serum truncated PA20 fragment. The intact mass of the isolated fragment was 23,600 Da. The predicted size of the fragment to SNSS is 23,588 Da, (thick arrow) a difference of 15 Da or 0.06 %, when compared with the measured mass. Underlined sequenceswere detected by MS analysis.The box represents the consensus recognition site for furin.

### Carboxypeptidase treatment of rPA

Given these results, we sought to determine whether this truncated PA_20_ fragment could result from serum carboxypeptidase digestion of PA_20_. Carboxypeptidases are a family of enzymes that cleave residues from the C-terminal end of a protein. This includes a group of enzymes that cleave basic amino acid residues from the carboxy terminus. To determine if carboxypeptidase could produce a truncated PA_20_ fragment, we conducted studies with a pancreatic carboxypeptidase. The effects of Carboxypeptidase B (CPB) treatment on furin-digested rPA were dose dependent. At higher concentrations (250 μg/ml; Figure 4 lane 5) multiple digestion fragments of PA were observed and PA_20_ reactivity was completely lost. A similar pattern was seen in the absence of furin and presumably relates to the presence of contaminating trypsin in this pancreatic preparation. In contrast at lower concentrations of CPB (25 μg/ml; Figure 4 lane 4), treatment produced a truncated PA_20_ fragment that was similar in size to that observed with serum digestion of PA (Figure 4, lane 1). Lower concentrations of CPB (2.5 μg/ml) had no effect on the size of furin-treated PA_20_, when compared with furin treatment alone.

**Figure 4.**
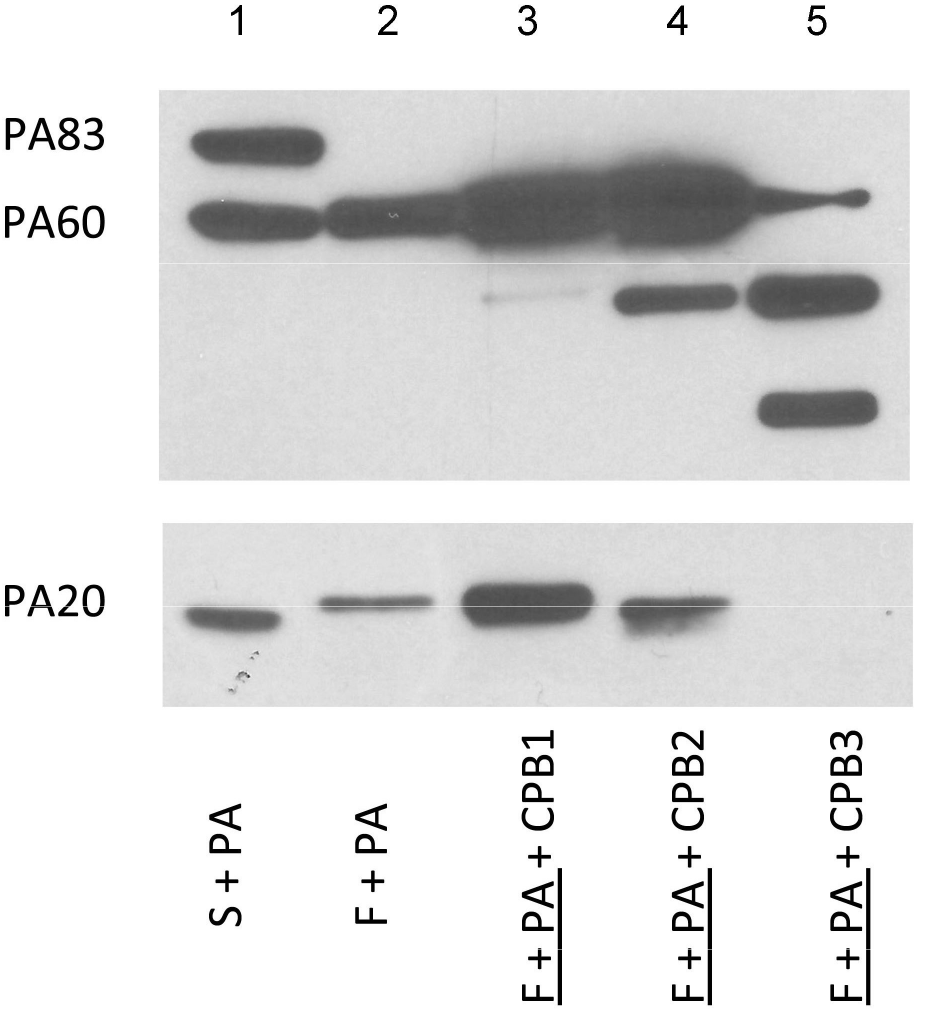
Carboxypeptidas B (COB) treatement of furn digested PA produces a truncated PA_20_ fragment. Treament of furin digested rPA83 with CPB from pig pancrease resulted ina dose related truncation of the PA_20_ fragment. This was most aparrent for CPB2 (25 *u*g/ml) as opposed to lower concentrations of CPB1 (2.5 *u*g/ml).Incubation with higher concentrations CPB3 (250*u*g/ml) resulted in complete loss of PA_20_ reactivity and apperence of multiple digestion fragments. Underline indicated pre-incubation

### Inhibition of serum carboxypeptidase activity

Next we sought to determine whether the ability of serum to produce a truncated PA_20_ fragment could be inhibited by carboxypeptidase inhibitors. Both Guanidinoethylmercaptosuccinic acid (GEMSA) potato tuber extract (PTI) are potent competitive inhibitors of carboxypeptidase though their inhibitory activity is not specific to any one class of carboxypeptidases. Addition of GEMSA, (500 μg/mL) to serum prevented the formation of a truncated PA_20_ and resulted in a PA_20_ fragment that was more similar in size to that produced by furin digestion (Figure 5). In contrast, no inhibition was seen with lower concentrations of GEMSA and for all concentrations of caboxypeptidase inhibitor (PTI) from potato-tuber extract.

**Figure 5.**
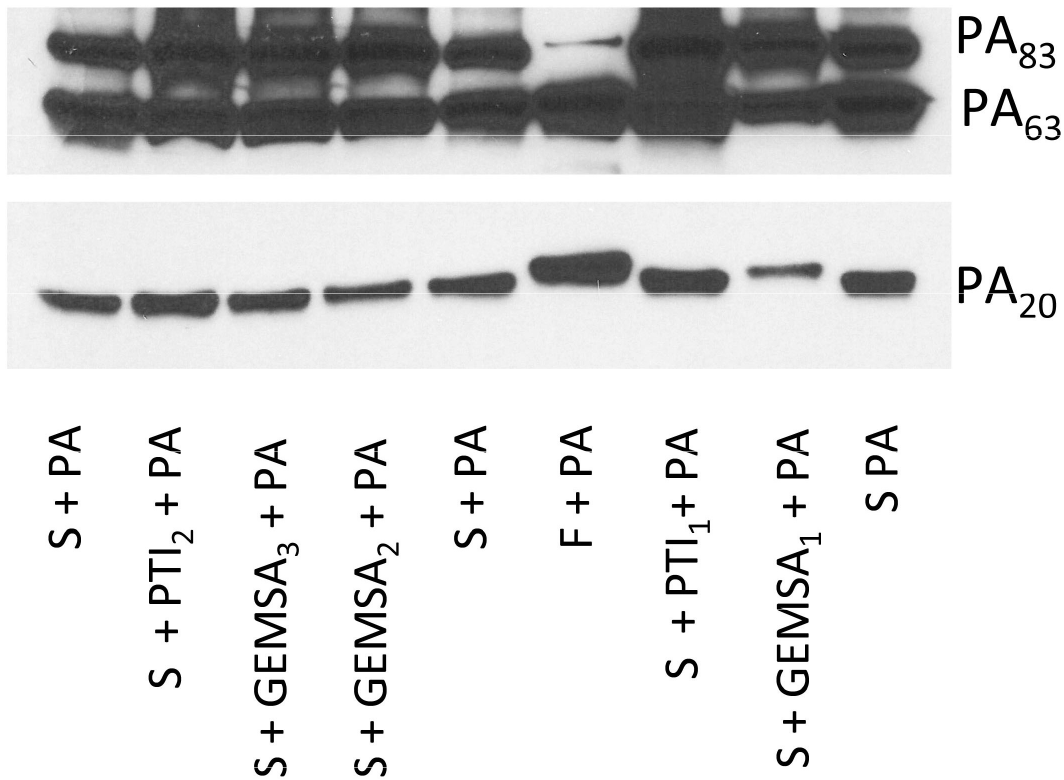
GEMSA but not caboxypeptidase inhibitor from potato-tuber extract (PTI) prevents formation of truncated PA_20_. In the presence of high concentrations of GEMSA (GEMSA-L, 500μg/ml), serum treatment of PA_83_ produced a PA_63_ fragment and a nontruncated PA_20_ fragment. In contrast, PTI at concentrations as high 1.25 mg/ml (PTI_1_) failed to inhibit serum truncation of PA_20_. Lower concentrations of GEMSA (50 and 5 μg/ml; GEMSA_2_and GEMSA_3_) and PTI_2_ (125 μg/ml) had no effect on serum-truncation of PA_20_.

## DISCUSSION

*B. anthracis* and the toxins it secrete have an intimate association with the circulation and serum over the course of infection. Our studies confirm earlier reports that both human and animal sera contain a furin-like enzyme, which digests PA to produce PA_63_ and PA_20_ fragments (13–15). In our own studies this activity was associated with protection against lethal toxin in vitro (15). We now extend these findings to demonstrate that human serum contains a carboxypeptidase, which further processes the PA_20_ fragment by removing the C-terminal basic amino acid residues, resulting in a truncated PA_20_ fragment. These findings contrast with the current model of anthrax toxin, which suggests that processing of PA occurs only at the cell surface and provide additional evidence for the complexity of anthrax toxin mechanisms of action. However, we note that serum and cell surface PA processing are not mutually exclusive events.

PA_20_ has been detected in the blood of *B. anthracis* infected animals though its contribution to anthrax pathogenesis is unknown (17). Nonetheless, several lines of evidence suggest may be play an active role in infection. For example, PA_20_ contains, a PA_14_ domain that is conserved among bacterial toxins and appears to play a role in cell binding (18).

Hammamieh et. al., reported that exposure of human peripheral blood mononuclear cells to PA_20_ induced a variety of genes related to the inflammatory, cell migration and triggered apoptosis in these cells (17). Furthermore, PA_20_ has been reported to bind Lethal Factor (19). Although circumstantial these findings are consistent with a role for PA_20_ in the pathogenesis of anthrax.

Serum is known to contain 2 carboxypeptidases (CP), CP-N and CPB2 (also known as CPU, plasma carboxypeptidase B and thrombin-activatable fibrinolysis inhibitor). Both carboxypeptidases cleave carboxy-terminal arginine and lysine residues from peptides/proteins and have been implicated in regulating inflammation through their actions on serum protein cascades, like the complement, anaphylatoxins, and kinins (20). As members of the carboxypeptidase family, both CP-N and CPB2 contain a zinc-binding site that makes them susceptible to inhibition by metal chelators. CP-N is constitutively produced by the liver with serum concentrations on the order of 30 μg/ml (21). In contrast, CPB_2_ must be activated by fibrin and once activated down-regulates fibrinolysis by removing terminal lysines from fibrin and is present in serum concentrations on the order of 4 − 15.0 μg/ml (22, 23). Elevated levels of CPB_2_ have been found in both animal models of bacterial sepsis and in septic patients and have been hypothesized to play a role in the hypercoagulability associated with sepsis (24–26). Interestingly, both carboxypeptidases inactivate complement anaphylatoxins (27, 28). Furthermore, both C3 and C5 have been implicated in the host response to anthrax (29, 30). Thus PA_20_ may possibly alter anthrax pathogenesis by interfering with anaphylatoxin inactivation during anthrax-associated sepsis.

It is interesting that CP-N is more susceptible to inhibition by GEMSA, while CPB2 is more susceptible to inhibition by potato carboxypeptidase inhibitor (31). Thus, our findings are consistent with the hypothesis that *in vitro,* CP-N is primarily responsible for the observed truncation of PA_20_. Nonetheless, the precise carboxypeptidase responsible for the truncation of PA_20_ *in vivo* (including during the sepsis of anthrax) is not known and it is likely that there is redundancy to the process. Of note, macrophages also express a membrane-associated carboxypeptidase (CP-M) that cleaves C-terminal lysines and arginine residues from proteins (32). It is, therefore, likely that a similar processing occurs at the surface of target cells.

In summary, we demonstrate that serum processing of PA is a 2-step process that involves a furin-like digestion of the PA_83_ component followed by truncation of the PA_20_ fragment by serum carboxypeptidases. The significance of these 2 serum-associated activities remains to be defined. Based out earlier studies that associate furin-like digestion with protection, we believe that this activity may in fact contribute to the host response to anthrax. This would be consistent with the close association of *B. anthracis* to the circulatory system. We also suggest that it is possible that the variation in these serum proteolytic activities contributes to differences in individual susceptibility to anthrax. Additional study looking and gain and lost of function in the context of experimental infection may help further delineate the important of these processes.

## MATERIALS AND METHODS

### PA

Recombinant PA_83_ (rPA) and its amino acid sequence were obtained from Wadsworth laboratories, New York State Department of Health (Albany, NY).

### Sera

Serum from lab volunteers was obtained and stored at −80°C with approval from the Committee of Clinical Investigations at Albert Einstein College of Medicine. In some experiments, pooled sera, processed to retain complement activity (Sigma, St Louis, MO) was used. These commercial sera produced comparable results to those obtained with sera from human volunteers.

### Antibodies and peptides

A library of 6 murine monoclonal antibodies (7.5G, 16A12, 10F4, 19D9, 20G7 and 2H9) that were previously generated and characterized was used to both define the digestion site and as detection reagents for immunoblot studies (33). Binding sites for these antibodies are provided in supplemental table 2. A previously synthesized library of overlapping peptides, which represents the PA sequence, was used for inhibition studies (16).

### Proteolytic Digestion and Fragment Detection

Proteolytic digestion studies were performed as previously described (15). Briefly, rPA (2.5 μg) was incubated with 25 μl of serum, phosphate buffered saline, or furin (0.5 Units, Invitrogen) at 37°C for 30 - 60 minutes. In some experiments, serum was heat-treated at 56 °C for 30 minutes prior to incubation with toxin. In other experiments, protease inhibitors (see below) or peptides at **a concentration of 5 μg/ml** were added to serum prior to incubation with rPA. Digested rPA was separated by SDS-electrophoresis and transferred to a nitrocellulose membrane. Membranes were blocked with 5% milk and then incubated with primary antibody. The following MAbs were used to characterize rPA cleavage: 10F4 (IgG1) and 7.5G (IgG2b). All MAbs were used at a concentration of 0.25 μg/ml. Primary antibody was detected with horseradish peroxidase-labeled goat isotype-specific antibody at a dilution of 1:25,000. Proteins were visualized by development with the ECL chemiluminescence kit (Pierce, Rockford, IL).

### Inhibition studies

#### Peptides

Serum (24 μl) was incubated with individual biotinylated peptides, peptide mixtures or PBS for 2 hours at room temperature. These peptides were chosen from a library of peptides representing the entire length of rPA and were synthesized as 15-mer, overlapping by 10 residues (16). This serum peptide mixture was then incubated with 1.5 μg of rPA for 30 min at 37° C and the resulting mixture subjected to separation by SDS page and detection by western blot

#### mAbs

PA (1.5 μg) was incubated with one of several PA-specific mAbs (2μg) (33) for 10 minutes at room temperature. This mixture was then added to 24 μl of serum, incubated at 37 °C for 20 minutes and then subjected to SDS electrophoresis and immunoblotting.

#### Protease inhibitors

A volume of 10 μl of sera was pre-incubated with 1 of 9 protease inhibitors included in a commercially available protease inhibitor set (Roche) for 30 minutes at 30° C. Individual inhibitors including (antipain, bestatin, chymostatin, E-64, phosphoramidon, pefabloc sc and aprotinin), each of which were reconstituted as per manufacturer’s instructions. Following this incubation 1.5 μg of rPA was added to the mixture and incubated at 37° C for 1 h. Specific inhibition of furin activity was accomplished using Furin inhibitor I (Caymen Chemicals) and Furin inhibitor II (Sigma). These compounds are selective competitive inhibitors of the proprotein convertases, including furin. Serum (12 μl) was incubated with furin inhibitors (at room temperature for 10 minutes and after which rPA (1.5 μg) was added and the entire mixture incubated for an additional 1 h at 37°C.

#### Carboxypeptidase inhibition

For these experiments sera was pre-treated with a variety of inhibitors for 30 minutes prior to incubation with rPA. These inhibitors included: guanidinoethylmercaptosuccinic acid (GEMSA, Santa Cruz Biotechnology), or carboxypeptidase inhibitor from potato tuber extract (Sigma). The serum PA digest mixture was separated by electrophoresis. PA_63_-like and truncated PA_20_ fragments were then detected with the antibodies 10F4 and 19D2 respectively.

#### Mass spectrometry (MS)

To isolate the truncated PA_20_ molecule, serum-digested rPA was incubated overnight at 4 °C with 200 μl of protein G resin in binding buffer (20 mM Tris, 150 mM NaCl, pH 7.4) together with the 50 μg of the mAb 19D2. The resultant slurry was centrifuged for 2.5 min at 2,500 G and the resin washed 5 times with binding buffer (Pierce). Following elution the protein was separated in a non-denaturing gel and electro-eluted for further analysis.

Mass spectrometric measurements (MS) and liquid chromatographic (LC) separations were obtained on the LTQ linear ion trap mass spectrometer (Thermo Scientific, San Jose, CA), the Rapid Separation LC 3000 (Dionex Corporation, Sunnyvale, CA) for tryptic peptides and the HP 1100 series for intact protein separation. For intact protein molecular weight measurements of the electro-eluted protein a C4 Vydac TP column (1 × 50 mm; 300 Å; 50 μL/min) was used. After desalting at 1% acetonitrile in 0.1% aqueous formic acid (FA) for 2 min the protein was eluted after increasing to 55% acetonitrile in 0.1% aqueous FA. The mass range from 600 to 1800 *m*/*z* was acquired on the LTQ and the raw data was deconvoluted using MagTran (34) or ProMass (ThermoFisher Scientific). Another aliquot of the electro-eluted protein was separated on a 1D SDS gel and selected molecular weight bands were excised for in-gel tryptic digestion as described (35). After sample injection and LC peptide separation (using an acetonitrile gradient), the top ten most abundant ions obtained from the survey scan (300 to 1600 *m/z)* were selected for fragmentation (MS/MS). Normalized collision energy of 35% and a 2 *m*/*z* isolation width were used for MS/MS. The MS/MS data were converted to a text file for peptide/protein identification using Mascot (Matrix Science Inc.).

#### Carboxypeptidase-mediated digestion of PA

To determine whether, carboxypeptidase digestion of furin treated rPA could produce a fragment similar in size to that seen with serum digestion of rPA, experiments were done with carboxypeptidase B (CPB) (Sigma). For these experiments, rPA was treated with furin for 10 minutes at 30 ° C and the mixture was incubated with CPB at different concentrations at 37° C. Proteins were separated by SDS PAGE and detected by immunoblotting as described above.

## ACKNOWLEDGMENTS

This work was supported, in whole or in part, by National Institutes of Health Grants AI33774-11, HL59842-07, AI33142-11, and AI52733-02 (to A. C.). This work was also supported by the Northeastern Biodefense Center under Grant U54-AI057158-Lipkin and the NIH-funded Shared instrumentation grant (1S10RR019352) for the LTQ LC-MS/MS system.

---

**Supplemental Table 1.**
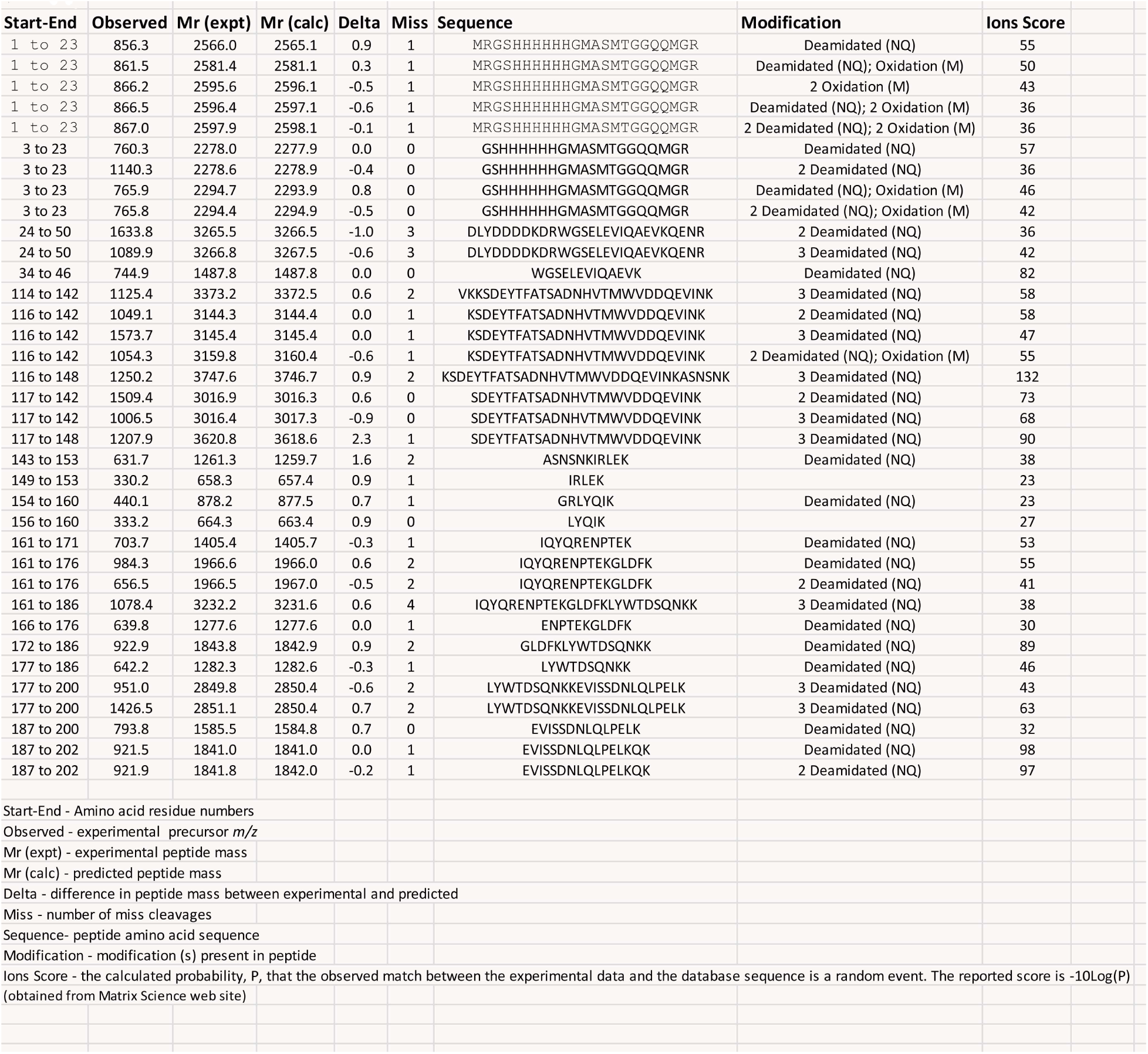

**Supplemental Table 2.**

| Antibody | Isotype | PA Fragment | Specificity |
| --- | --- | --- | --- |
| 7.5g | IgG<sub>2b</sub> | PA<sub>20</sub> | Domain 1 |
| 10F4 | IgG<sub>1</sub> | PA<sub>63</sub> | Domain 4 |
| 19D9 | IgG<sub>1</sub> | PA<sub>20</sub> | Domain1* |
| 20G7 | IgM | PA<sub>20</sub> | Domain 1* |
| 2H9 | IgG<sub>1</sub> | PA<sub>63</sub> | Domains 2-4 |

These antibodies compete with each other to bind Domain 1.

**Supplemental Figure 1.**
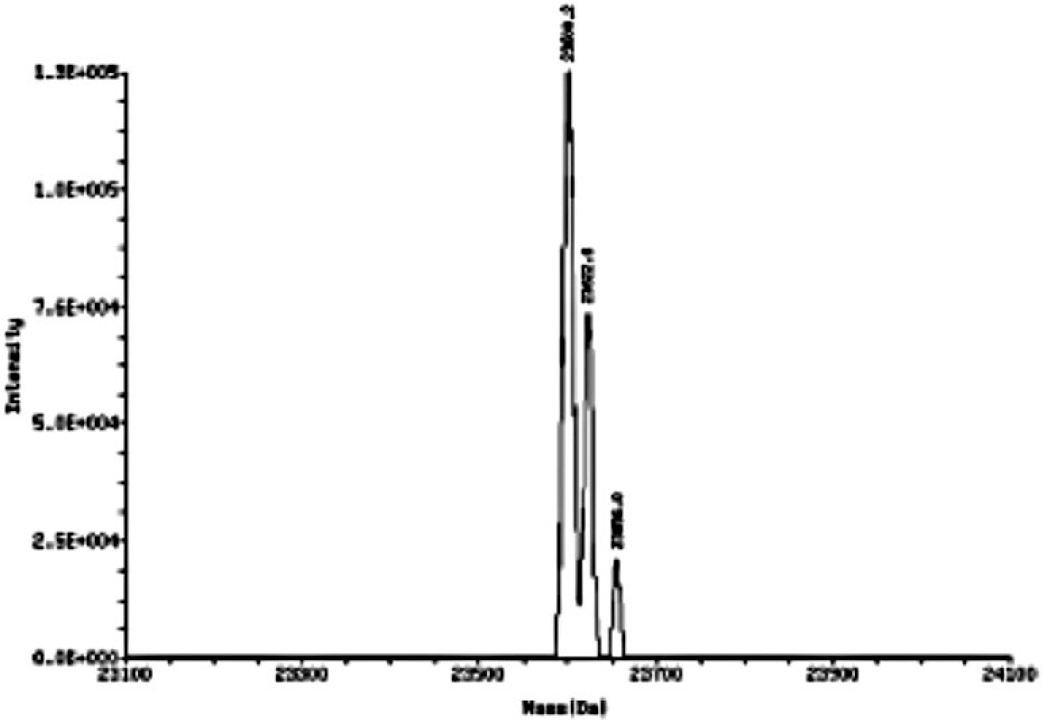
Deconvoluted experimental mass for the truncated PA20 fragment obtained from the intact protein LC-MS measurement.

**Supplemental Figure 2.**
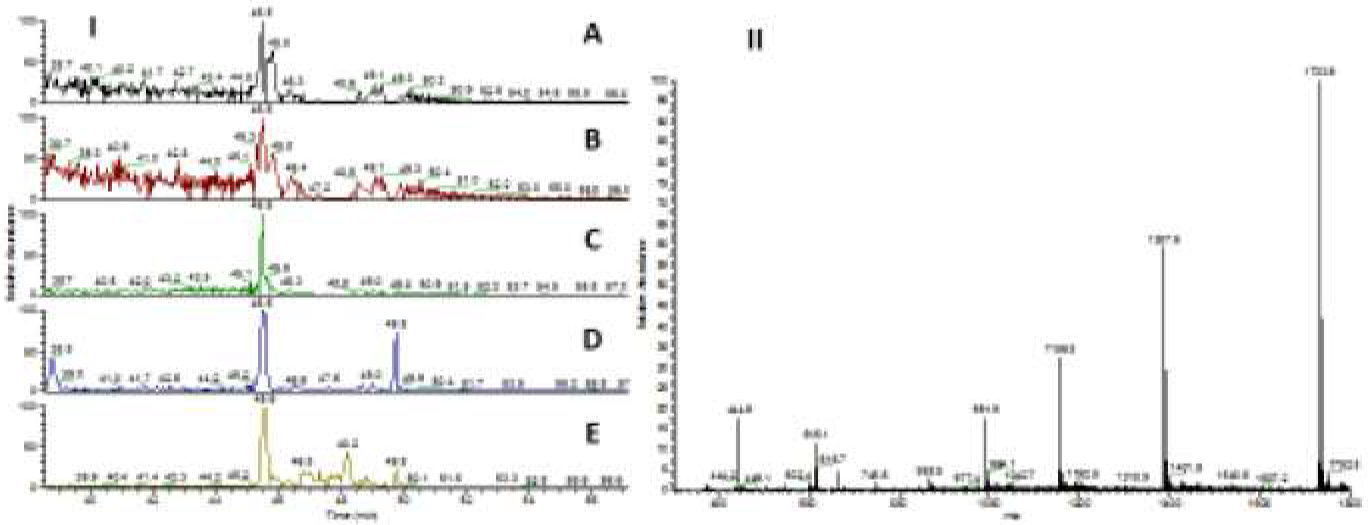
Extracted ion chromatogram (I)ofthe +4, +5, +6 + 7, +8 charge states (A-E) of the peptide rPA83 following trypsin digestion-LLNESESSSQGLLGYYFSDLNFQAPMWTSSTTGDLSIPSSELNIPSE NQYFQSAIWSGFIK. The masss spectra for the 45.6 minute retention time peak corresponding to this peptide is shown(II).

